# Design and Development of Point of Care Test and Optical Reader for Early Screening of Kidney Related Disorder

**DOI:** 10.1101/2020.07.01.181024

**Authors:** Pankaj Shihvare, Satyam Mohla, Tejal Dube, Alok Verma, Rohit Srivastava

## Abstract

Low-cost, paper-based colorimetric assays for early screening of albumin, creatinine and their ratio have been developed. The developed methods are noninvasive and require only 10µl of the urine sample. A reflectance-based optical reader has also been developed for the quantification of the albumin and creatinine. The developed method is based on spot urine testing which is advantageous when compared to the conventional 24-hour urine collection. The detection range of albumin and creatinine assays is 10-150 mg/dl and 25–400 mg/dl, respectively. The developed assays and optical reader were tested with the chronic kidney diseased patient’s samples at KEM Hospital, Mumbai.

## I. Introduction

In many high-income countries, the prevalence of chronic kidney disease approaches 15% of the adult population, predominantly due to the large diabetic population and hypertensive nephropathy[1, 2, 3, 4, 5]. In India, there are more than 10 million cases every year for chronic kidney disease (CKD), which is a long-term form of kidney disease affecting 5–10% of the Indian population[6, 7, 8, 9]. Unfortunately, the symptoms of worsening kidney function are not specific, and lack of early symptoms makes the diagnosis difficult.

Preeclampsia is another life-threatening disorder that occurs during pregnancy, childbirth and the postpartum period. Proteinuria is one of the major features of preeclampsia, which is the most common cause of complications during pregnancy. Hypertensive disorders of pregnancy (which include preeclampsia) are one of the most common causes of death due to pregnancy[10, 11, 12]. Women who have had preeclampsia are at increased risk of heart disease and stroke later in life[13, 14]. Protein and/or albumin to creatinine ratio are shown to have a most prominent biomarker for early detection of CKD and preeclampsia[15, 16]. The current methods to detect elevated albumin level in urine are a lab test or a dipstick assay. In resource-limited countries like India, almost 65% of the population lives in rural and suburban areas and do not have access to good medical facilities. Thus, the current method to detect albumin and creatinine levels is only the standard lab test, which requires sophisticated instruments and trained lab technicians. Majority of Indian population is undiagnosed to pre-diabetes and diabetes, which is the major cause of renal related disorders. Many reports suggest that morbidity, mortality and economic burden from renal-related disorders are likely to be a growing public health concern for India[7, 8]. Apart from that in critical cases, the clinician needs to track the albumin and creatinine ratio many times in a day which is very difficult for individual patients. If one needs to measure the random albumin to creatinine ratio, there are several colorimetric strips available in the market, but most of them are expensive, which make them unaffordable for the mass population.

Test for quantification of urinary albumin excretion in terms of albumin to creatinine ratio can be performed to predict albuminuria accurately and becomes increasingly relevant in assessing the prognosis and treatment of kidney diseases. Degree of albuminuria reflects the progression of kidney diseases which helps in assessing the prognosis of kidney diseases affecting the normal kidneys functions, and if no treatment is given in time, it could lead to severe complications including kidney failure[15, 16].

Estimation of albumin to creatinine ratio in urine is used for not only monitoring but also for treatment purposes. Traditionally, the assessment of albuminuria is done by collecting a 24-hr urine sample to measure the amount of albumin excreted in mg per 24-hr. This method is fairly accurate but becomes unreliable due to the problem of storage environment of the urinary samples, sedimentation of the proteins and ions. Also, the collection and storage of the urinary samples for 24-hr are too tedious for the patients and clinicians[17, 18]. For the past many years, the investigators have started collecting short term urinary samples such as 2-hours, 4-hr and spot-urine samples instead of tedious 24-hr urine collection. They have expressed the results in term of albumin to creatinine ratio measurements on a single-voided specimen provide a convenient and reliable alternative method than 24-hr urine measurements[19, 20, 21, 22].

Urine dipstick assays are an easy and simple alternative for analyses of albumin level, but their high cost is the main barrier. Apart from that, the urine dipstick assays give only the qualitative assessment of albumin level, which in some of the cases is not sufficient[21, 22, 23, 24, 25]. With this, we propose lowcost dye-based colorimetric dipstick assays for albumin and creatinine and a point-of-care optical reader for the quantitative assessment of the assays.

The development of low-cost albumin and creatinine assays can help a majority of the population to early diagnose kidney-related disorders such as CKD, pre-eclampsia. The developed albumin to creatinine ratio (ACR) measurement is an easy and simple alternative to the lab test where the patient has to collect the urine for 24-hr. As many researchers have reported that on-spot urine ACR has a high correlation with 24-hr urine albumin excretion, the developed low-cost assay will certainly help the patients by reducing the turnaround time of the test.

## II. Materials and Methods

Polypropylene glycol 400, bromophenol blue (BPB), cumene hydroperoxide, hydroxypropyl methylcellulose, 3, 3’, 5, 5’-tetramethylbenzidine (TMB), glycerol 2 phosphate were procured from Sigma Aldrich, USA. Citric acid, sodium citrate, copper sulphate, sodium chloride, sodium hydroxide, ethanol, bovine serum albumin, creatinine was purchased from Sigma Aldrich, USA. Other common laboratory reagents were obtained from SpectroChem Pvt. Ltd. All solvents and chemicals used in the experiments were of analytical grade. They were used as received without any further alteration and purification.

The immunoassay paper (absorbent pad) for colorimetric assay were procured from GE Healthcare. Different grades of immunoassay papers (CF4, CF3, CF1, 3mm CHR, and Chromatography) were used to get optimized dye-binding capacity and to have the maximum range for albumin and creatinine concentration due to color change. The CF4, CF3 and CF1 absorbent pads differ by their thickness, CF4 was the thickest with a thickness almost 400 µm. CF3 and CF1 were having a thickness of 250 µm and 150 µm, respectively.

### A. A. Preparation of Colorimetric Assay for Albumin

#### Binder

The polypropylene glycol was used as a binder to immobilize the dye onto the absorbent pad. 10polymer binder was prepared in the milli-Q. A small sheet of absorbent paper (Grade CF3) was cut and dipped into the polymer binder solution for 5 minutes and after that kept in the oven for drying at 50°C for 45 minutes. The template is used to format your paper and style the text. All margins, column widths, line spaces, and text fonts are prescribed; please do not alter them. You may note peculiarities. For example, the head margin in this template measures proportionately more than is customary. This measurement and others are deliberate, using specifications that anticipate your paper as one part of the entire proceedings, and not as an independent document. Please do not revise any of the current designations

#### Citrate buffer

Bromophenol blue is also used as the pH indicator dye. Thus, we need to stabilize the pH of the developed assay to diminish any effect of the acidic or basic pH in the colorimetric assay. Citrate buffer of 0.1M and 3.0 pH was prepared in the Milli-Q to minimize the effect of the urine pH. The absorbent pad was dipped into the 0.1M citrate buffer solution for 5 minutes and kept in the oven for drying at 50°C for 45 minutes.

#### Dye preparation

50 µg/ml of bromophenol blue dye was prepared in 99dipped into the prepared dye solution for 5 minutes, and then the excess dye solution was recovered, followed by drying in a hot air oven at 45°C for 30 minutes. The developed assay strip was then stored at room temperature for further experiments and testing

### B. Optimization of Colorimetric Assay for Albumin

Various concentrations of the polymer binder were used to optimize the BPB dye-binding capacity onto the paper substrate and prevent leaching of the dye in case of the excess urine sample. 0.5M, 0.1M, 1.5M, 2M of citrate buffer with different pH (2.5, 2.7, 3.0, 3.3) was used to optimize and diminish any effect of the urine pH on the developed assay. Different concentrations of BPB dye were tried to get optimized color change and detection range with the bovine serum albumin (BSA). Fig.1 shows the developed paper based colorimetric assay for the albumin. From the Fig.1, it is clear that the detection range of the developed assay is from 10-150 mg/dL, after 150 mg/dL the color change of the assay is not significant and gives almost the same results. It is because of the limitation of the dye-binding capacity of the immunoassay paper. For albumin concentration greater than 150 mg/dL, the samples can be diluted, and after testing it can be recalculated for the original concentrations.

**Fig. 1.**
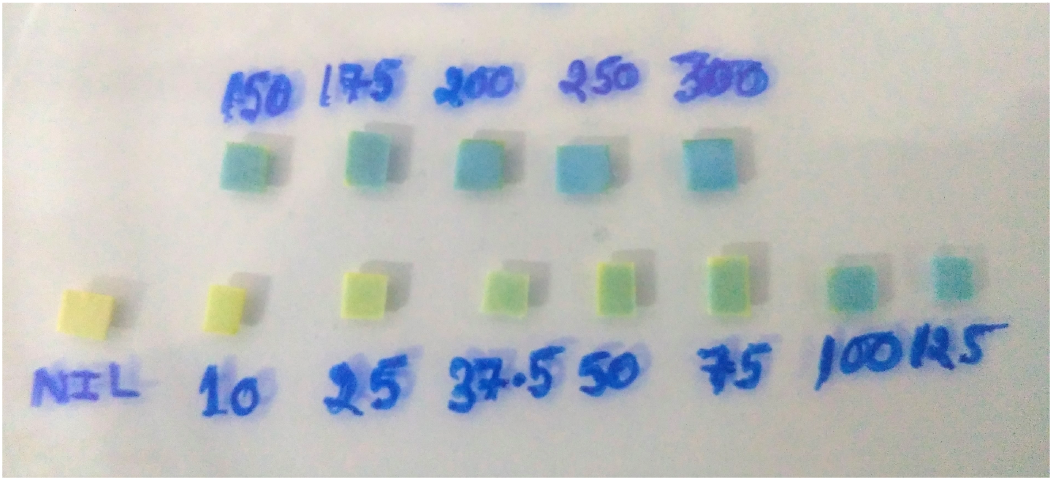
Developed colorimetric assay for albumin. Bovine serum albumin (BSA) was used as standard and assay was tested using the different concentration (mg/dL) of the BSA stock solution.

### C. Preparation of Colorimetric Assay for Creatinine

#### Binder

In creatinine assay, hydroxypropyl methylcellulose (HPMC) was used as a binder to immobilize the dye onto the absorbent pad. 1the Milli-Q. A small sheet of absorbent paper (Grade CF3) was cut and dipped into the polymer binder solution and kept in the oven for drying at 50°C for 50 minutes.

#### Copper citrate complex

CuSO4 salt was used as the source of the cupric ions and citric acid was used for the formation of copper citrate complex so that copper should not bind to the TMB. NaCl was used to stabilize the copper creatinine complex. 16 mM of CuSO4, 27 mM of citric acid, 500 mM of NaCl and 1 M of glycerol 2 phosphate was added in 10 ml of Milli-Q to prepare glycerol phosphate buffer. Glycerol 2 phosphate buffer provides the essential condition for the oxidation of the TMB in the presence of creatinine and cumene hydroperoxide. The pH of the above solution was adjusted to the 6.8 using 10M NaOH. The absorbent pad was dipped into the copper citrate buffer solution for 5 minutes and kept in the oven for drying at 50°C for 50 minutes.

#### Dye preparation

80 mM of TMB dye and 90 mM or 167µl of cumene hydroperoxide was prepared in 9 ml of absolute ethanol and 1 ml of acetonitrile. The TMB dye is photosensitive, so all the work was carried out in a dark room. The absorbent pad was dipped into the TMB dye solution for 5 minutes, and then the excess dye solution was recovered, followed by drying in room temperature 30°C for 90 minutes.

### D. Optimization of Colorimetric Assay for Creatinine

Paper-based creatinine assay was optimized with the similar method used with albumin assay. various concentration of the HPMC was used to optimize the TMB dye-binding capacity with the paper substrate and prevent leaching of the dye in case of the excess sample. Glycerol phosphate buffer with different pH (6.5 6.8, 7.0, 7.4) was used to diminish any effect of the urine pH on the developed assay. Various concentration of TMB dye was tried to get optimize color change and the maximum range with the creatinine standard. A small quantity of acetonitrile was used to increase the solubility of the TMB in ethanol. Fig.2 shows the developed paper-based colorimetric assay for the creatinine. From the Fig.2, it is clear that the detection range of the developed assay is from 25-400 mg/dL. Creatinine is a waste product produced by muscles from the breakdown of creatine, and the normal range of creatinine in the spot urine sample is 300-400 mg/dL.

**Fig. 2.**
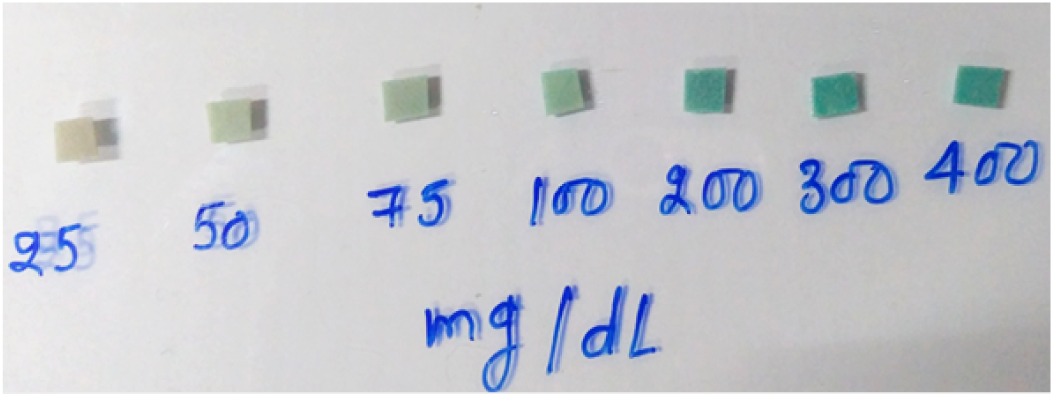
Developed colorimetric assay for creatinine. Creatinine was used as standard and assay was tested using the different concentration (mg/dL) of the creatinine stock solution.

## III. DEVELOPMENT OF OPTICAL READER

A reflectance-based optical reader was developed for the quantification of the paper-based albumin and creatinine assays, as shown in Fig.11. The development of optical reader was divided into three categories; (a) Light illumination module, (b) Reflectance sensing module, (c) Electronic hardware. The light illumination module consists of an array of LEDs to ensure uniform light radiation over the assay strip. The radiation wavelength and angle of the LEDs were selected based on the excitation wavelength of the dyes used for the assays. The bromophenol blue (BPB) and TMB excitation wavelengths are 605 nm and 650 nm, respectively. As shown in Fig.3, LEDs with different wavelengths were used for the assays.

**Fig. 3.**
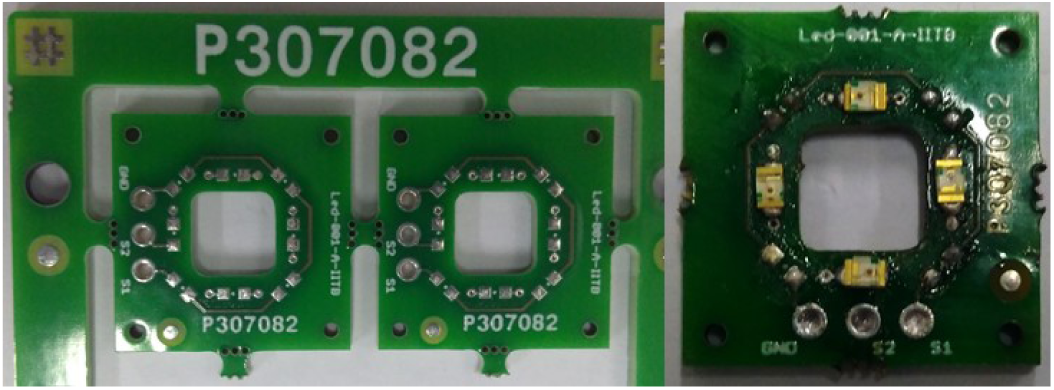
Developed colorimetric assay for creatinine. Creatinine was used as standard and assay was tested using the different concentration (mg/dL) of the creatinine stock solution.

**Fig. 4.**
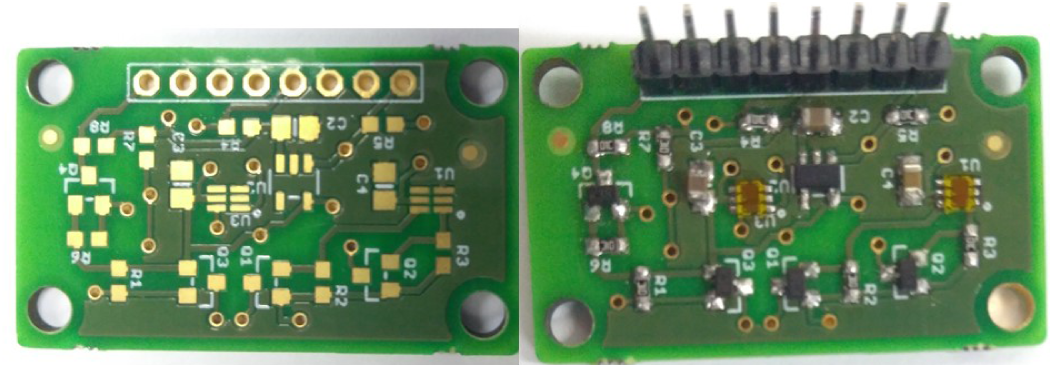
Digital Light photodiode with 95 % normalized spectral response in the range of 600-650 nm.

**Fig. 5.**
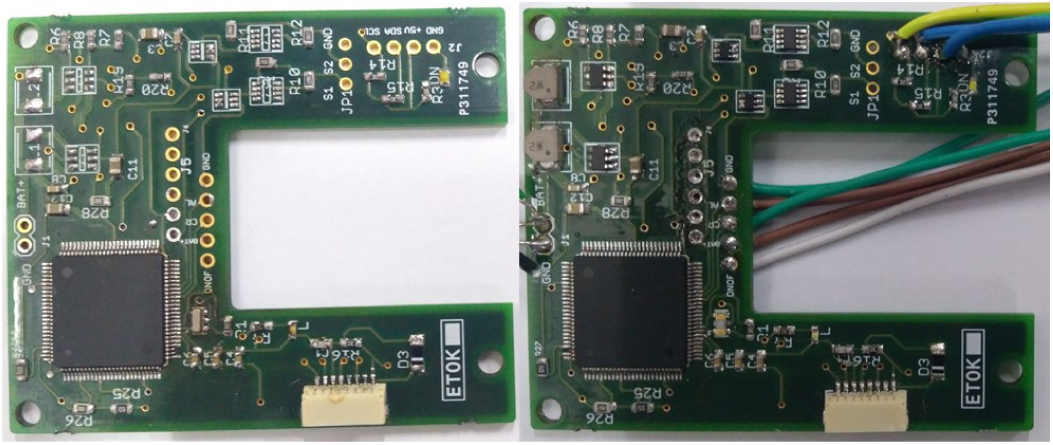
Fabricated PCB to handle interrupts and to perform all the instructions of the point-of-care reader operation.

**Fig. 6.**
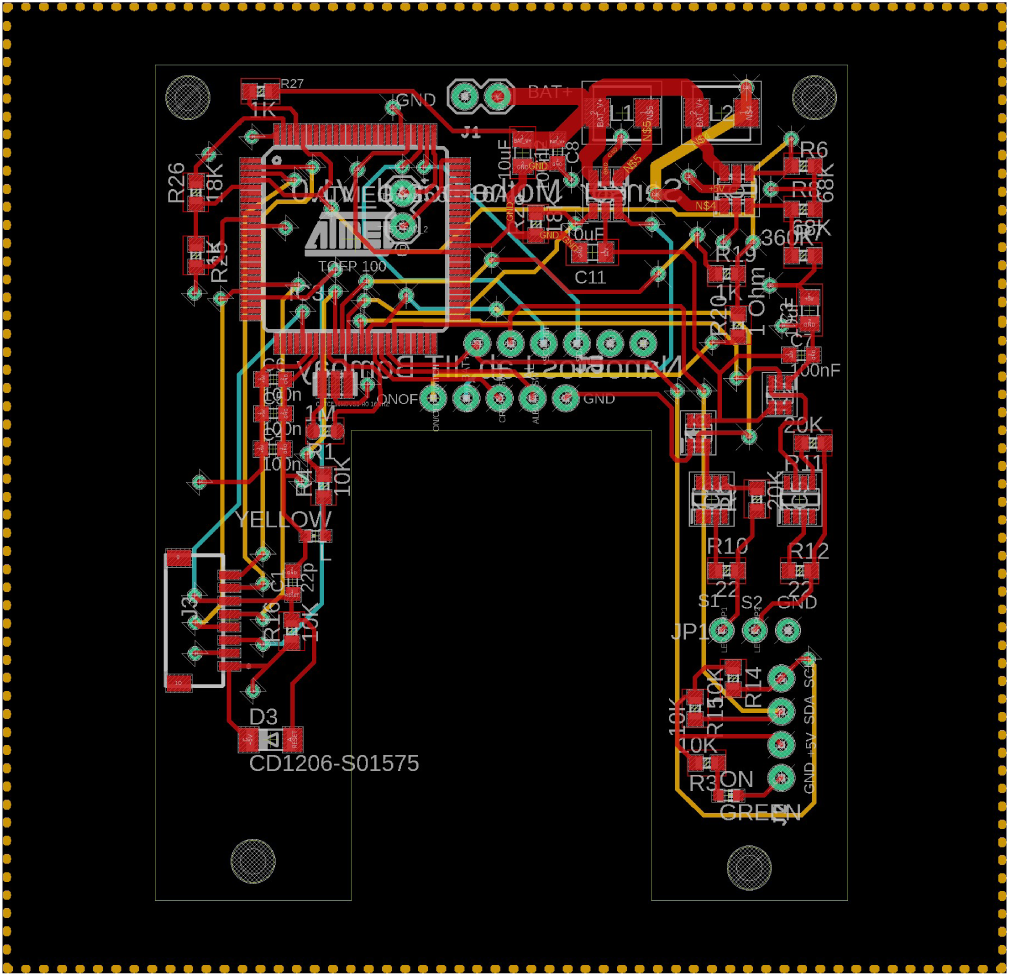
Proposed hardware architecture for the development of point-of-care reader for the colorimetric assay.

**Fig. 7.**
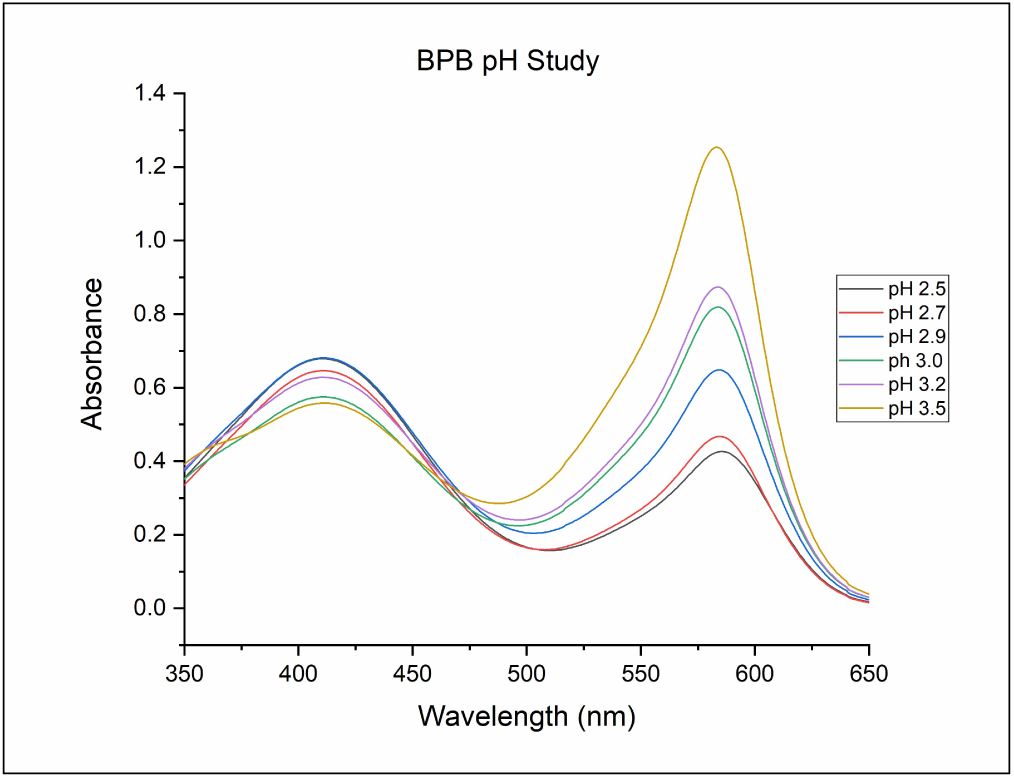
Effect of citrate solution with different pH in the absence of bovine serum albumin.

**Fig. 8.**
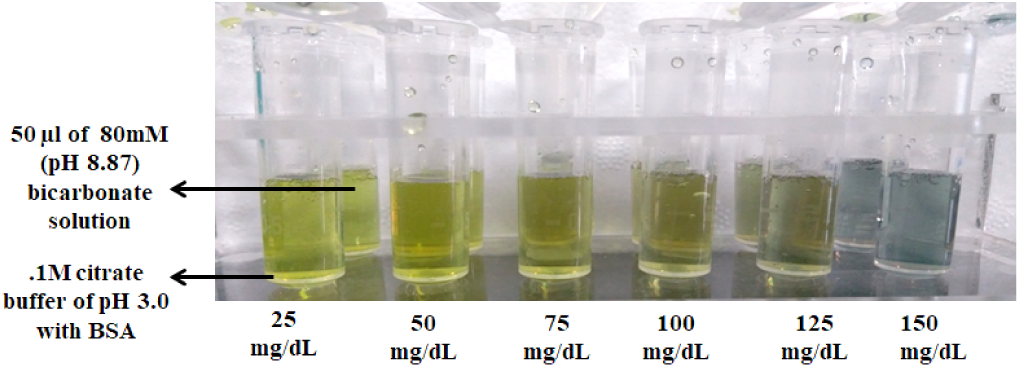
pH and molarity of the citrate buffer was optimized to diminish any effect (color change due to pH) on the albumin assay. Citrate buffer of 0.1M and 3.0 pH have almost no effect of alkaline urine samples and does not give any change in color due to alkaline pH of the urine sample.

**Fig. 9.**
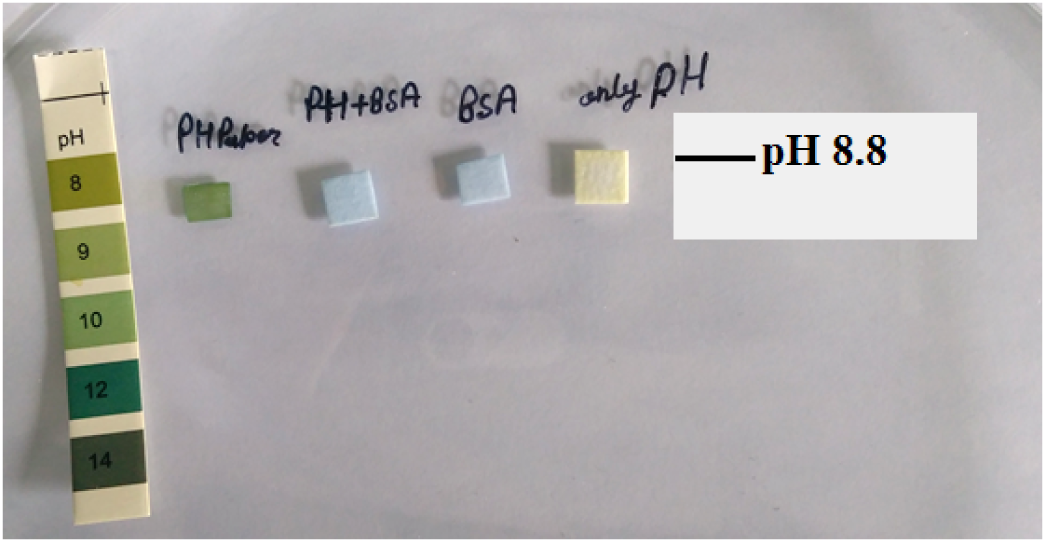
Paper-based assay with citrate buffer of 0.1M and 3.0 pH have almost no effect of alkaline urine samples and does not give any change in the color due to alkaline pH of the urine sample.

**Fig. 10.**
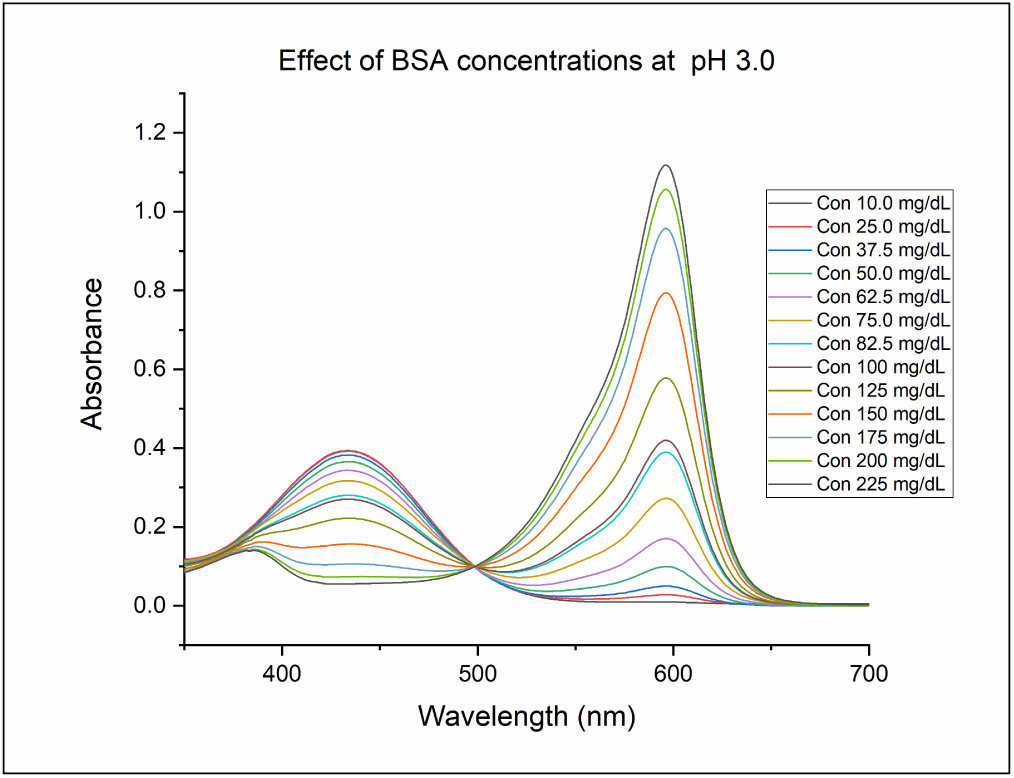
Interaction of bromophenol blue with different concentrations of bovine serum albumin at citrate buffer pH 3.0.

**Fig. 11.**
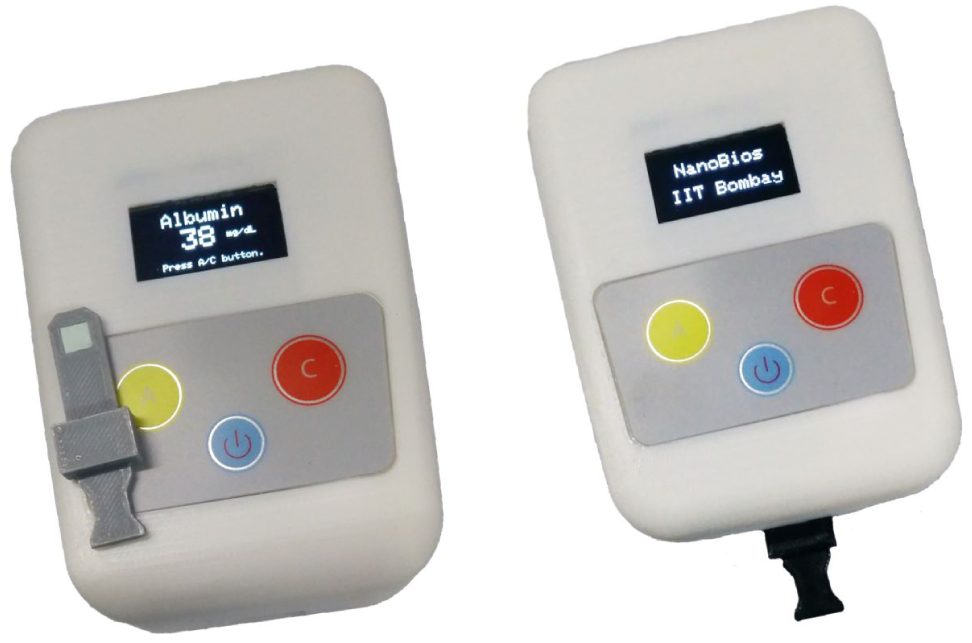
Developed reflectance-based point-of-care optical reader for the quantification of the albumin and creatinine assays.

The reflected light from the assays was quantified with the help of a photodiode. The photodiode was selected based on the maximum spectral sensitivity at the emission range of the BPB and TMB dye, which is 600-650 nm. The selected photodiode is a highly sensitive digital light sensor and has integrated light to digital value converter. The photodiode has an ultraviolet light filter, and it is selective to visible light only. It transforms light intensity into a digital signal output and capable of the direct I2C interface. The high sensitivity of the photodiode enables to detect the smallest change in the fluorescence intensity, as shown in Fig.4. The electronic hardware was designed using Autodesk eagle freeware printed circuit board (PCB) designing software. Atmel 2560 microcontroller was used as it has the high number of I/O pins, serial and I2C interface. Fig.5 and 6 shows the schematic architecture and fabricated PCB of the developed hardware board. 1.8-inch TFT screen and SD-card card were integrated to display and store the results for analysis purpose. The software to execute and control all the instruction of the reader was designed using the open-source Arduino integrated development environment.

## IV. RESULTS AND DISCUSSIONS

BPB dye is widely used in the pH indicator papers and changes its color as the pH of the solution changes. The effect of the pH of the urine solution was studied for the albumin assay, to find the optimized pH and molarity of the citrate buffer solution for the development of the assay.

### A. Effect of pH of the Urine on the Albumin Assay

The effect of the pH of the urine sample on the albumin assay was studied. Normally the pH of the urine is slightly acidic but certain conditions such as chronic kidney disease, urinary tract infection, acute kidney infection etc. can cause a change in the pH of the urine from acidic to alkaline.

0.05M citrate buffer solutions of different pH were prepared, and each solution was mixed with 50 µl of BPB solution. The absorbance spectra of each solution were taken with the help of Perkin Elmer UV-Vis spectrophotometer and shown in Fig.7. It showed that BPB dye reacted with the pH of the solution and gave the color change, which was not desirable for the proposed albumin assay.

To minimize the effect of pH of the urine sample, the molarity of the citrate buffer was increased from 0.05 M to 0.1 M. To mimic the pH of the alkaline urine, 80 mM bicarbonate solution of pH 8.9 was prepared in Milli-Q and tested with different concentration of the bovine serum albumin with 0.1M citrate buffer of pH 3.0. As shown in Fig.8 and 9, the bicarbonate of pH 8.9 solution does not affect the solution, and the color appearance (because of the interaction of BPB and BSA) with and without bicarbonate solution is the same. The paper-based albumin assay was then prepared with the 0.1M citrate buffer of pH 3.0 and tested with bicarbonate solution of pH 8.9.

As seen in Fig.9, the pH of the bicarbonate solution does not affect the developed assay. As seen in Fig.7, the absorbance peak of the bromophenol blue is at 580 nm, but the Fig.10 shows that, in the presence of bovine serum albumin, the absorbance peak shifted to 598 nm. The peak of the solution shifted from 580 nm to 598 nm due to the electrostatic interaction between bromophenol blue and bovine serum albumin[19].

Different concentrations of the BSA were prepared using serial dilution and tested with the above-prepared solution, and the absorbance spectra of each solution were taken with the help of Perkin Elmer UV-Vis spectrophotometer and shown in Fig.10. Fig. 10 showed that there was a visible difference among the different concentrations of the BSA, which can be detected with the help of the point-of-care reader.

### B. Testing and Validation of the Developed Assays and Optical Reader

The developed assays for albumin and creatinine were tested with different concentration of albumin and creatinine standards in the lab as well the chronic kidney diseased patient’s urine samples with the collaboration of KEM hospital, Mumbai, India as shown in Fig.12 and 13. BSA and creatinine standard were used for lab testing. Serial dilutions were prepared with different concentrations, and 10 µl of the standard solution was used as the sample. Fig. 12 and 13 shows that optical reader was reproducible and sensitive enough to differentiate among the various concentrations of albumin and creatinine.

**Fig. 12.**
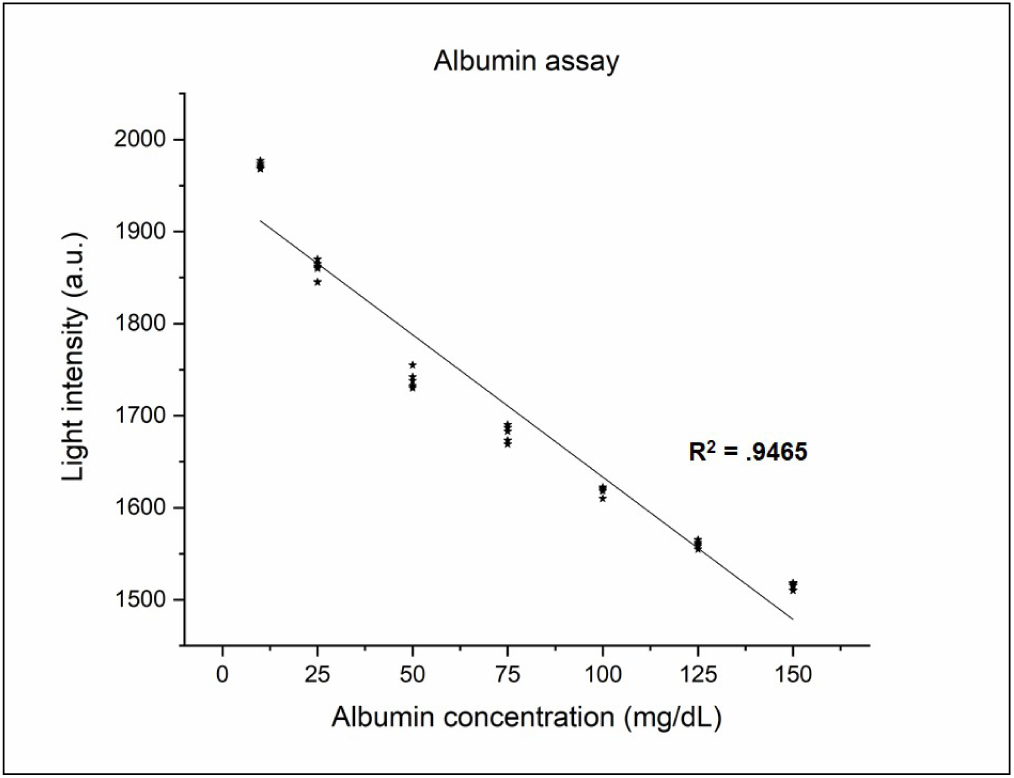
Interaction of bromophenol blue with different concentrations of bovine serum albumin at citrate buffer pH 3.0.

**Fig. 13.**
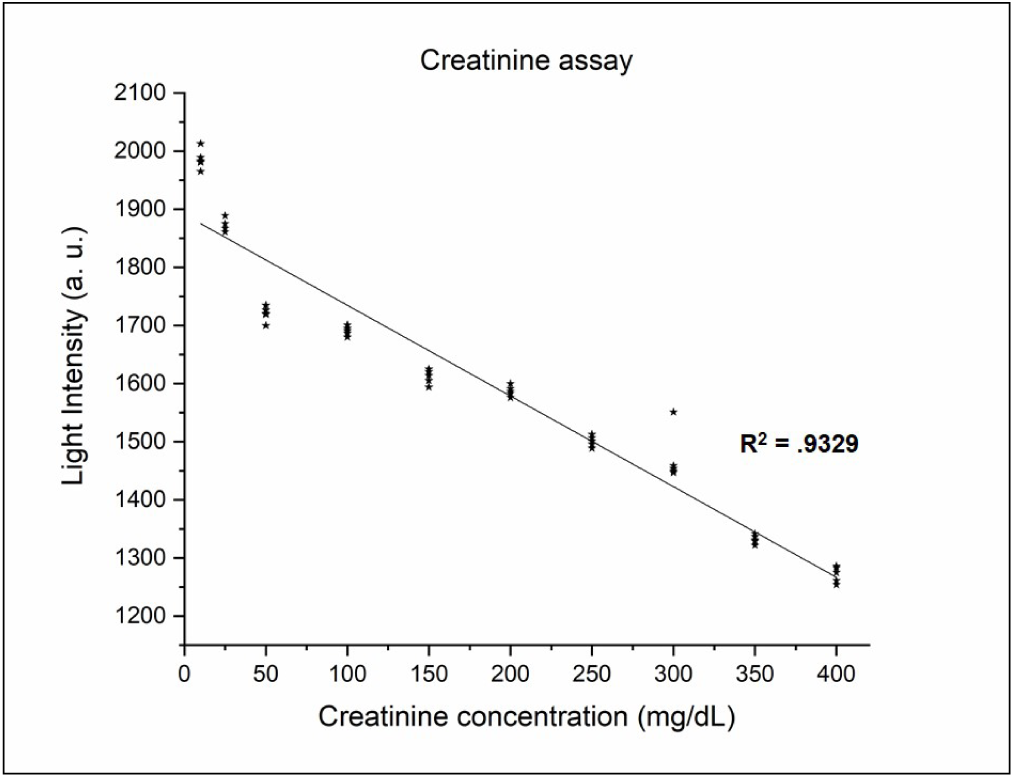
Testing of the paper-based creatinine assay with creatinine standards with different concentrations.

## V. CONCLUSION

Paper-based colorimetric assay for albumin, creatinine and point-of-care optical reader for the quantification of the same were successfully developed. The developed assays were low-cost as dyes were used instead of enzymes or antibodies attached to gold nanoparticles to indicate the color change when reacted with the analytes. The albumin and creatinine assays were optimized by using different concentrations of the polymer binder, molarities of the buffer, various pH of the buffers and dyes concentrations. Effect of the urine pH was studied on the albumin assay as BPB dye is used as a pH indicator. By increasing the molarity of the citrate buffer, the color change due pH of the urine solution was diminished. The testing of the assays and optical reader was done using albumin and creatinine standards as well as the patient’s samples at KEM hospital, Mumbai, India. The detection range of the albumin assay is 10-150 mg/dL and for creatinine is 25-400 mg/dL. The optical reader was reproducible and sensitive to differentiate the various concentrations of albumin and creatinine when tested. The results were found to be close in agreement.

## ACKNOWLEDGMENT

The present research is supported by Department of Biotechnology (DBT), India. All authors acknowledge the support from Dr Sucheta Dandekar and Ms Radhika Mhatre for the clinical validation of the assays and optical reader at KEM hospital, Parel, Mumbai, India.

## Notes

### Competing Interest Statement

The authors have declared no competing interest.

## References

[1] Josef Coresh et al. “Prevalence of chronic kidney disease in the United States”. In: Jama 298.17 (2007), pp. 2038–2047.

[2] Carl A Burtis and David E Bruns. Tietz fundamentals of clinical chemistry and molecular diagnostics-e-book. Elsevier Health Sciences, 2014.

[3] Marion Kerr et al. “Estimating the financial cost of chronic kidney disease to the NHS in England”. In: Nephrology Dialysis Transplantation 27.suppl 3 (2012), pp. iii73–iii80.

[4] Luxia Zhang et al. “Prevalence of chronic kidney disease in China: a cross-sectional survey”. In: The Lancet 379.9818 (2012), pp. 815–822.

[5] LA Kaplan, AJ Pesce, and SC Kazmierczak. “Clinical chemistry: theory, analysis, correlation 3rd ed”. In: Maryland Heights: Mosby.[Google Scholar] (1996).

[6] Anna J Dare et al. “Renal failure deaths and their risk factors in India 2001–13: nationally representative estimates from the Million Death Study”. In: The Lancet Global Health 5.1 (2017), e89–e95.

[7] P Devi et al. “Prevalence, risk factors and awareness of hypertension in India: a systematic review”. In: Journal of human hypertension 27.5 (2013), pp. 281–287.

[8] Vivekanand Jha and Gopesh Modi. “Uncovering the rising kidney failure deaths in India”. In: The Lancet Global Health 5.1 (2017), e14–e15.

[9] Sreejith Parameswaran et al. “Referral pattern of patients with end-stage renal disease at a public sector hospital and its impact on outcome”. In: National Medical Journal of India 24.4 (2011), p. 208.

[10] Amir Said Alizadeh Naderi and Robert F Reilly. “Primary care approach to proteinuria”. In: The Journal of the American Board of Family Medicine 21.6 (2008), pp. 569–574.

[11] Elosha Eiland, Chike Nzerue, and Marquetta Faulkner. “Preeclampsia 2012”. In: Journal of pregnancy 2012 (2012).

[12] H Wang et al. “Global, regional, and national life expectancy, all-cause mortality, and cause-specific mortality for 249 causes of death, 1980â 2015: a systematic analysis for the Global Burden of Disease Study 2015”. In: The Lancet 388.10053 (2016), pp. 1459–1544.

[13] Noura Al-Jameil et al. “A brief overview of preeclampsia”. In: Journal of clinical medicine research 6.1 (2014), p. 1.

[14] M Gojnic et al. “Plasma albumin level as an indicator of severity of preeclampsia.” In: Clinical and experimental obstetrics & gynecology 31.3 (2004), p. 209.

[15] Baha Sibai, Gus Dekker, and Michael Kupferminc. “Pre-eclampsia”. In: The Lancet 365.9461 (2005), pp. 785–799.

[16] Joseph A Abdelmalek et al. “Estimated albumin excretion rate versus urine albumin-creatinine ratio for the assessment of albuminuria: a diagnostic test study from the Prevention of Renal and Vascular Endstage Disease (PREVEND) Study”. In: American Journal of Kidney Diseases 63.3 (2014), pp. 415–421.

[17] ACG Collins et al. “An aid to the early detection and management of diabetic nephropathy: assessment of a new point of care microalbuminuria system in the diabetic clinic”. In: Diabetic medicine 18.11 (2001), pp. 928–932.

[18] Kouri et al. “Evaluation of the ARKRAY AUTION Eleven® reflectometer in detecting microalbuminuria with AUTION Screen® test strips and proteinuria with AUTION Sticks 10PA® strips”. In: Scandinavian Journal of Clinical and Laboratory Investigation 69.1 (2009), pp. 52–64.

[19] Yong-ju Wei, Ke-an Li, and Shen-yang Tong. “The interaction of bromophenol blue with proteins in acidic solution”. In: Talanta 43.1 (1996), pp. 1–10.

[20] J Sereikaite, Z Bumeliene, and V Bumelis. “Bovine serum albumin-dye binding”. In: Acta Chromatographica 15 (2005), p. 298.

[21] Anna Michnik, Katarzyna Michalik, and Zofia Drzazga. “Stability of bovine serum albumin at different pH”. In: Journal of Thermal Analysis and Calorimetry 80.2 (2005), pp. 399–406.

[22] Thomas W Stephens. Assay for the determination of creatinine. US Patent 4,529,708. 1985.

[23] Michael J Pugia. Oxidative creatinine assay. US Patent 5,374,561. 1994.

[24] Kin F Yip, Amy H Chu, and Brenda Tudor. “Method for the determination of urinary protein and creatinine”. In: Biotechnology Advances 4.13 (1995), p. 785.

[25] Sabine Rode et al. “Complexation chemistry in copper plating from citrate baths”. In: Journal of the Electrochemical Society 151.6 (2004), p. C405.

